# Monocyte microRNA signatures in late-onset neonatal sepsis

**DOI:** 10.64898/2026.10.15.759337

**Authors:** Eleni Gkouti, Adamantia Kouvela, Eleni Papachatzi, Maria Apostolidi, Gabriel Dimitriou, Vassiliki Stamatopoulou, Despoina Gkentzi

## Abstract

**Purpose:** Neonatal late-onset sepsis (LOS) remains a major cause of neonatal morbidity and its non-specific presentation challenges early diagnosis. MicroRNAs (miRNAs) are promising biomarkers, but unbiased next-generation sequencing (NGS) of monocyte-specific miRNA has not been applied to neonatal LOS. We aimed to identify the monocyte-specific miRNA expression profile in neonatal LOS and its associations with the neutrophil-to-lymphocyte ratio (NLR) and systemic immune-inflammation index (SII).

**Methods:** In a prospective study at the University General Hospital of Patras, 30 samples from 15 neonates with LOS and 8 healthy controls were analyzed. Monocyte miRNA expression was characterized by NGS pre and post antimicrobial therapy in cases and controls. Predicted target genes of the major candidate miRNAs were evaluated by RT-qPCR in an independent cohort. Clinical severity was assessed using NLR and SII.

**Results:** Hsa-miR-6743-3p was significantly upregulated on day 0 and returned to control levels by day 2. Four additional miRNAs (hsa-miR-7108-5p, hsa-miR-3940-5p, hsa-miR-6851-3p and hsa-miR-1909-3p) exhibited a consistent trend toward upregulation in acute LOS. NLR, but not SII, was significantly elevated at diagnosis and showed excellent diagnostic performance (AUC 0.958), normalizing by day 2. Hsa-miR-7108-5p correlated positively with NLR on day 0. In the RT-qPCR cohort, its predicted target *DRAM2*, was reduced on day 0 and partially recovered by day 2.

**Conclusion:** Monocyte miRNA profiling identified transient alterations in neonatal LOS that parallel clinical severity, while NLR showed high discriminatory performance. These exploratory findings warrant validation in larger cohorts to verify monocyte miRNAs as potential biomarkers of neonatal LOS.

**What is Known:**

- LOS diagnosis is challenging, as blood cultures take 36–72 hours to yield results
- NLR and SII may complement established biomarkers, although findings vary
- Circulating miRNAs are promising biomarkers, but no consistent signature has been identified

**What is New:**

- Unbiased NGS identified alterations in monocyte miRNA expression in neonatal LOS
- hsa-miR-6743-3p was upregulated in acute LOS and returned to control levels by day 2
- Additional miRNAs showed a consistent trend toward upregulation during acute LOS

## Introduction

Neonatal late-onset sepsis (LOS) remains a major cause of morbidity and mortality in neonatal intensive care units (NICUs) [1]. Early diagnosis is challenging due to nonspecific clinical signs, while blood culture, the diagnostic gold standard, requires at least 36–72 hours for definitive results [1].

Inflammatory biomarkers such as C-reactive protein (CRP) and procalcitonin (PCT) are widely used in neonatal sepsis; however, neither provides sufficient diagnostic accuracy alone, prompting interest in combined biomarkers [2]. Moreover, traditional markers provide limited information on disease severity and treatment response [2], increasing interest in accessible, low-cost complete blood count-derived indices, particularly neutrophil-to-lymphocyte ratio (NLR) and systemic immune-inflammation index (SII) [3]. Recent studies reported higher NLR/SII values in neonatal sepsis (than controls), with SII showing the strongest discriminatory ability [3]; while NLR was independently associated with LOS, showing high diagnostic performance (AUC of 0.903) [4]. However, findings remain inconsistent among studies [5] and therefore current evidence supports NLR and SII as complementary rather than stand-alone diagnostic markers.

MicroRNAs (miRNAs) are small, single-stranded non-coding RNAs, that regulate gene expression post-transcriptionally. Since their discovery in 1993 [6], over 2,000 human miRNAs have been identified. They regulate diverse biological processes, including development, differentiation, metabolism, and immune responses [7], and are estimated to target over 60% of human protein-coding genes [8]. MiRNA dysregulation has been implicated in cancer, cardiovascular disease, and sepsis, highlighting their potential as diagnostic and therapeutic targets [9]. The stability of circulating miRNAs makes them promising biomarkers for neonatal sepsis with pooled sensitivity and specificity of 83% and 76%, respectively [10]. To present, several miRNAs are differentially expressed in septic neonates. MiR-34a-5p and miR-199a-3p are downregulated and correlate with SNAP-II severity scores [11], miR-16 is upregulated with better diagnostic performance than CRP/PCT, whereas miR-181a and miR-23b are downregulated [12]. Evidence remains limited and a recent scoping review identified only 12 LOS-specific studies and no consistent miRNA signature [1]. However, previous work used targeted RT-qPCR of predefined miRNA candidates in serum or plasma rather than comprehensive NGS-based miRNA profiling of isolated monocytes. The aim of the present study was therefore to characterize the miRNA expression profile of monocytes in LOS using NGS and to determine whether differential miRNA expression was associated with clinical severity, as reflected by NLR and SII. Finally, predicted target genes were evaluated by RT-qPCR in an independent group of neonates with clinical LOS.

## Materials and methods

### Population and clinical data

Neonates admitted to the NICU at the University General Hospital of Patras who developed clinical LOS at ≥72 hours of life and received antimicrobial treatment for at least 10 days were matched with healthy controls from the same unit with no evidence of sepsis. In total, 38 samples were analyzed: 30 samples from 15 neonates with LOS, collected on day 0 (before antibiotic administration) and day 2 (48 hours post-antibiotics), and 8 samples from controls. Clinical LOS was supported by clinical features (Online Resource 1) and laboratory findings, including the sepsis score. Exclusion criteria were current weight <500 g, necrotizing enterocolitis at any time, congenital infections, and any surgical procedure. Cases and controls were matched for gestational age, sex, birth weight, race, day of life, parity, and socioeconomic background. Socioeconomic status was estimated using residential postal codes as a proxy measure. Demographic and clinical data were obtained from medical records (12/2023-12/2025). To assess cohort representativeness, the demographic characteristics of enrolled neonates were compared with all neonates who developed LOS during the same period; no significant differences were identified (data not shown). Patients and controls were anonymized and labeled as follows:

NGS cohort:

– Patients 01 to 10 (P01–P10): LOS cases (culture-positive)
– Controls 01 to 06 (C01–C06): matched controls

RT-qPCR cohort:

– Patient 11 (P11): LOS case (culture-positive sepsis)
– Patients 12 to 15 (P12–P15): LOS cases (culture-negative clinical sepsis)
– Controls 07 to 08 (C07–C08): matched controls

### Systemic inflammatory indices

NLR and SII were calculated from complete blood counts obtained at each blood draw (day 0, 2) using absolute cell counts (×10 /L). NLR was calculated as the absolute neutrophil count divided by the absolute lymphocyte count, and SII (N×PLT/L) as the absolute neutrophil count × platelet count / absolute lymphocyte count. Both indices were compared between days 0 and 2 and with healthy controls and analyzed in relation to differential miRNA expression.

### Blood sampling and laboratory analyses

Following parental written informed consent, 1–2 mL of peripheral blood was collected from cases (day 0 and 2) and controls. Mononuclear cells were isolated by Lymphoprep density-gradient centrifugation (2000 rpm, 18°C, 20 min) and stored in RNAlater (Invitrogen) at −80°C. Total RNA was extracted using the mirVana miRNA Isolation Kit (Invitrogen) and quantified using the Qubit RNA HS Assay Kit. miRNA-enriched cDNA libraries were prepared using the Ion Total RNA-Seq Kit v2 and sequenced on an Ion GeneStudio S5. Reads were mapped to mature human miRNAs from miRBase v22 using Bowtie2 (v2.4.2) [13], quantified using featureCounts (v2.0.1), and analyzed with limma-voom (v3.50.1) [14,15]. Differential expression was assessed between controls and day 0, day 0 and day 2, and controls and day 2, including miRNAs detected in all three groups. Significance was defined as log2FC >0.5 and p<0.05; p=0.05–0.10 indicated suggestive trends. Sequencing data are available in GEO.

Predicted targets of differentially expressed miRNAs, identified using miRDB [16,17] were evaluated by RT-qPCR in an independent cohort. RNA was extracted from isolated monocytes and reverse-transcribed using M-MuLV Reverse Transcriptase (NEB) and random hexamers. Target genes were quantified using gene-specific primers with ACTB as the reference gene (Online Resource 2). Statistical comparisons between day 0 and day 2 were performed using a paired t-test.

### Statistical analysis

Continuous variables are reported as median (IQR) and categorical variables as n (%). LOS cases and controls were compared (days 0, 2) using the Mann–Whitney U test, and paired day 0–2 values using the Wilcoxon signed-rank test. Systemic inflammatory indices (NLR, SII) are presented separately for the NGS and RT-qPCR cohorts, while comparisons were performed on pooled cases and controls. The diagnostic accuracy of day 0 NLR was assessed by ROC analysis (AUC with 95% CI; optimal cut-off by Youden index). NGS-identified miRNAs differing at day 0 (log2FC >0.5, p<0.05) or showing a suggestive trend (p=0.05–0.10) were correlated with NLR and SII at days 0 and 2 within the NGS cohort (P01–P10) using Spearman’s rank correlation (ρ).

#### AI Use Disclosure

During the preparation of this manuscript, the authors used Claude (Anthropic), a large language model-based AI tool, to assist in the creation of the figures. All AI-assisted content was critically reviewed and verified by the authors, who take full responsibility for the accuracy and integrity of the manuscript.

## Results

### Demographics

Demographic and clinical characteristics of both cohorts, including gestational age, birth weight, sex, day of life at sepsis onset, causative pathogen, respiratory and inotropic support, and the presence of central venous lines are summarized in Table 1. The total study sample included 15 LOS cases and 8 controls, with median gestational age 34.6 weeks (IQR 32.0–35.7) vs 34.0 weeks (IQR 32.1–38.2) and median birthweight 2140 g (IQR 1665–2668) vs 2165 g (IQR 1760–2635), respectively.

**Table 1.**
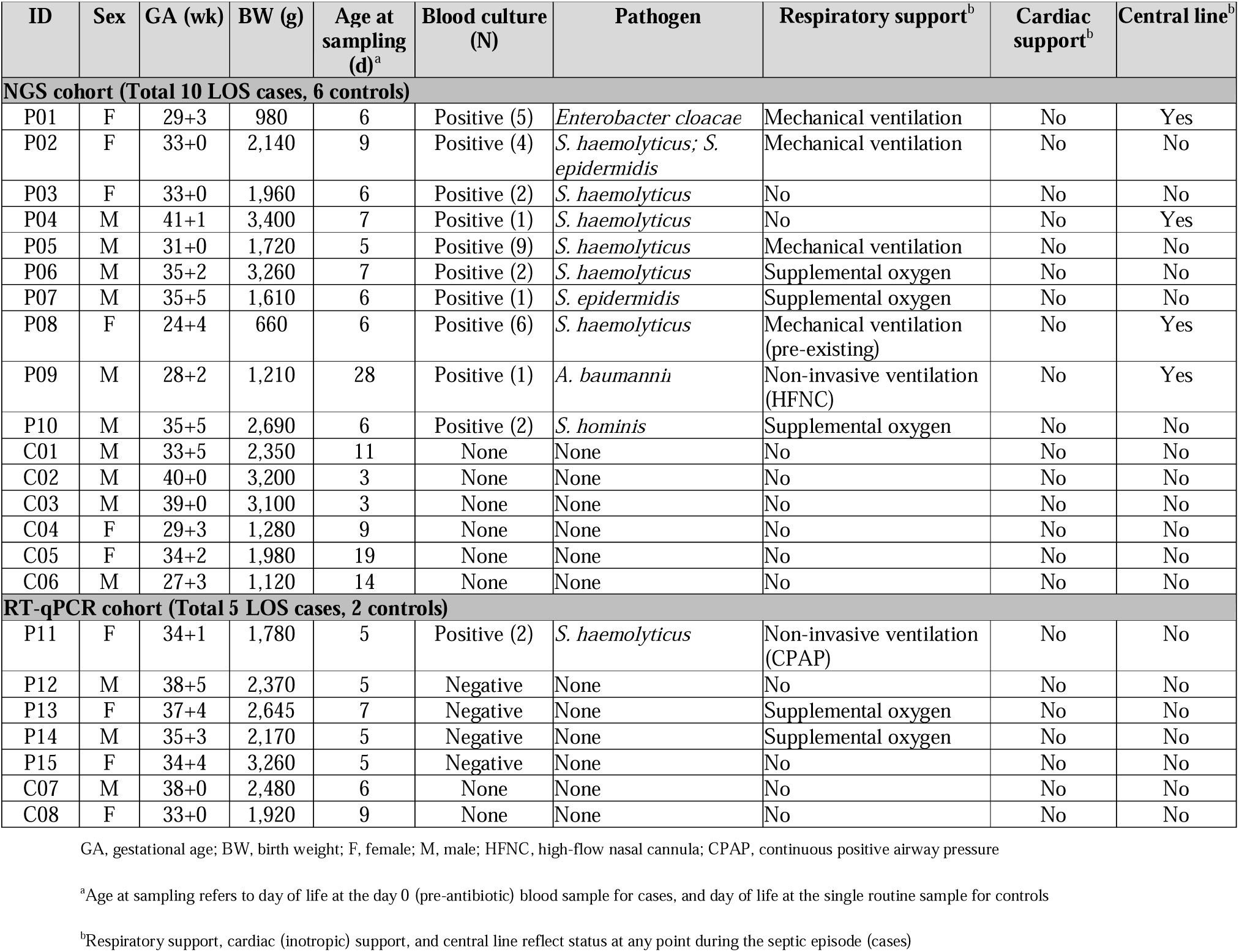
Demographics, clinical characteristics and pathogen causing LOS in the study cohorts.

### Systemic inflammatory indices

To support the clinical diagnosis of LOS in the study cohort, NLR and SII values were compared between LOS cases (n=15) on days 0 and 2 and controls (n=8). At day 0, NLR was significantly higher in LOS cases compared to controls (median 5.49 [IQR 2.48–9.11] vs. 1.10 [IQR 0.53–1.80]; p=0.0001). SII followed a similar directional pattern but did not reach statistical significance (p=0.132). Both indices decreased significantly between day 0 and day 2 (NLR: p=0.0001; SII: p=0.0003), and neither differed significantly from controls at day 2 (NLR: p=0.796; SII: p=0.190), consistent with clinical recovery after initiation of antimicrobial therapy (Figure 1).

**Fig. 1.**
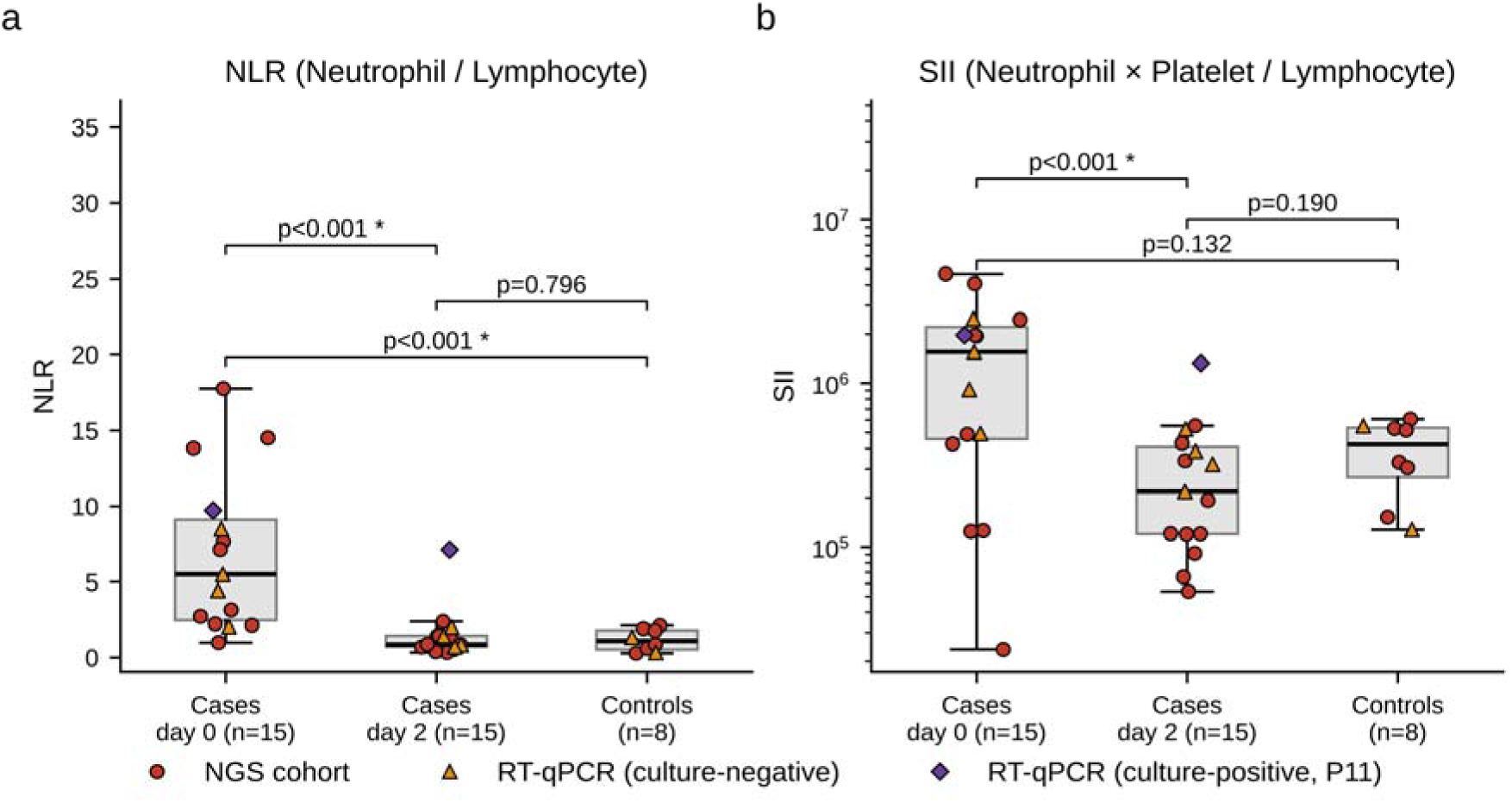
Systemic inflammatory indices (NLR and SII) in neonates with late-onset sepsis (LOS) at diagnosis (day 0) and 48 hours post antibiotics (day 2), compared with healthy controls (LOS cases, n = 15; healthy controls, n = 8). (a) Neutrophil-to-lymphocyte ratio (NLR). (b) Systemic immune-inflammation index (SII). Circles denote the NGS cohort, triangles the culture-negative RT-PCR cohort cases, and the diamond the culture-positive RT-PCR cohort case. Boxes show the median and interquartile range; whiskers extend to the most extreme values within 1.5 × the interquartile range. Brackets show p-values (Mann–Whitney U test for cases vs controls; Wilcoxon signed-rank test for day 0 vs day 2); an asterisk (*) indicates p < 0.05

Findings were consistent when the NGS and RT-qPCR cohorts were analyzed separately (Online Resource 3). In the NGS cohort, NLR was significantly higher in cases than controls at day 0 (median 5.14 [IQR 2.35–12.29] vs. 1.31 [0.67–1.88]; p=0.0017), decreased significantly by day 2 (p=0.002), and did not differ significantly from controls at day 2 (p=0.664). SII was numerically higher in cases than controls at day 0 (median 1,220,384 [IQR 201,779–2,330,000] vs. 424,638 [312,116–528,430]; p=0.562) and decreased significantly by day 2 (p=0.0098). In the RT-qPCR cohort, both NLR and SII were numerically higher in cases than controls at day 0 (NLR: median 5.49 [IQR 4.44–8.50] vs. 0.84 [0.58–1.09], p=0.095; SII: median 1,560,000 [IQR 916,588–1,973,160] vs. 339,596 [233,755–445,436], p=0.191) and decreased from day 0 to day 2 in all five cases (both p=0.0625). Given its significant discrimination between LOS cases and controls, ROC analysis was performed for NLR on day 0. The marker (NLR) demonstrated strong diagnostic potential for identifying LOS cases within our test group (AUC 0.958, 95% CI 0.867-1.000) and was used in the present study as a benchmark for LOS cases in the subsequent microRNA analysis (Online Resource 4).

Small RNA sequencing was performed in the NGS cohort comprising 10 neonates with blood culture-confirmed LOS and 6 matched controls. For the LOS cases, samples collected on day 0 and 2 (n=20 samples) were analyzed separately, as described above. Following sequencing and bioinformatic analyses, 33, 38, and 34 mature human miRNAs were identified in the controls, day 0, and day 2 samples, respectively (Figure 2a). Out of them 29 miRNAs were consistently detected in all three study groups and included in the subsequent differential expression analysis.

**Fig. 2.**
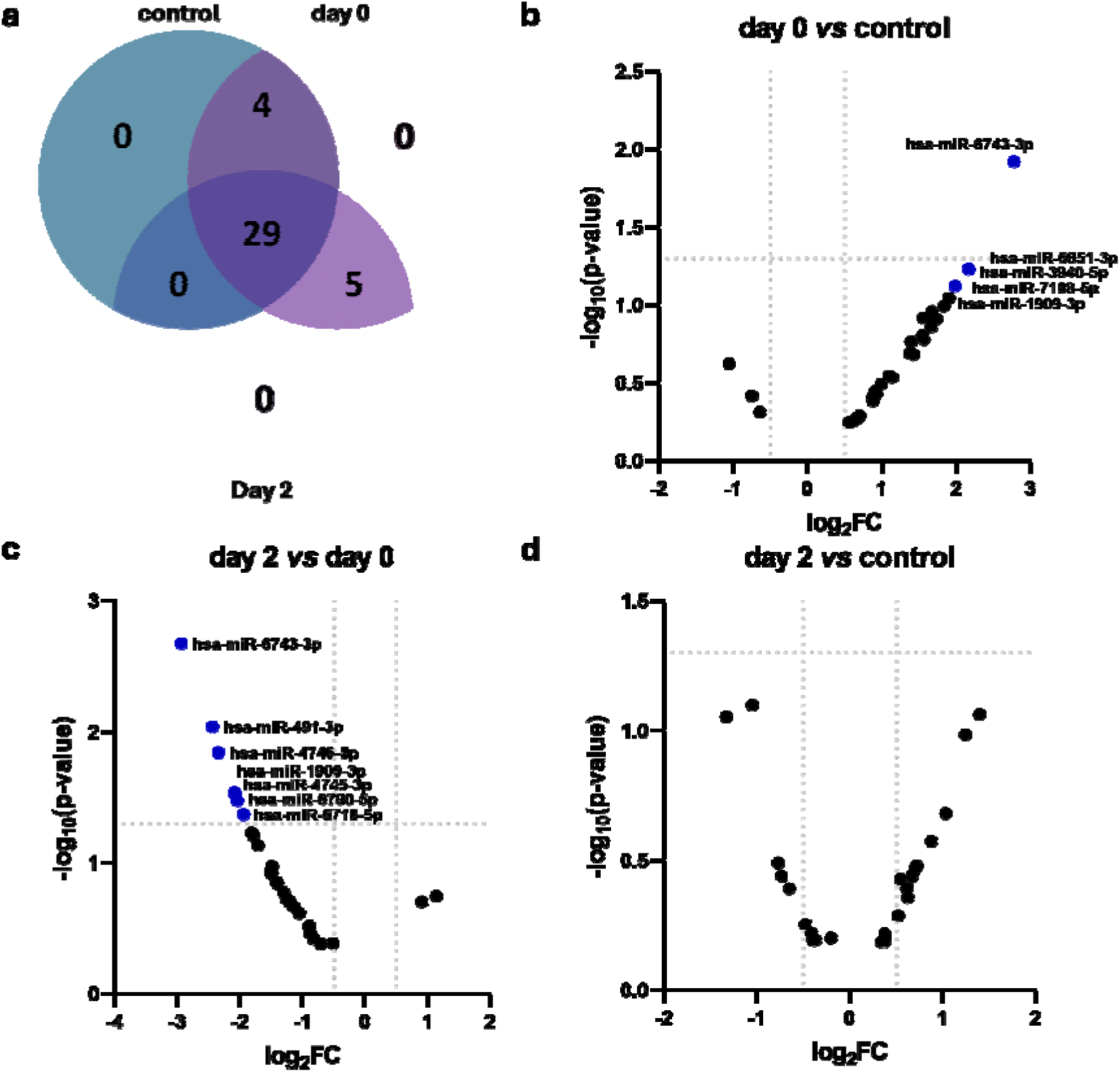
Differential miRNA expression profiles in neonatal monocytes at the diagnosis of late-onset sepsis (LOS) and 48 hours post antibiotics (NGS cohort; LOS cases, n = 10; healthy controls, n = 6). (a) Venn diagram showing the overlap of detected miRNAs among healthy controls, neonates with LOS at diagnosis (day 0), and 48 hours post antibiotics (day 2). (b–d) Volcano plots of differential miRNA expression for day 0 versus control (b), day 2 versus day 0 (c), and day 2 versus control (d). The x-axis represents log fold change (log FC) and the y-axis −log (p-value). Vertical dotted lines indicate the fold-change thresholds (log FC = −0.5 and 0.5), and the horizontal dotted line indicates the significance threshold (p = 0.05)

At day 0, hsa-miR-6743-3p was significantly upregulated in neonates with LOS compared to control samples (Figure 2b). Five additional miRNAs, hsa-miR-7108-5p, hsa-miR-3940-5p, hsa-miR-6851-3p and hsa-miR-1909-3p, showed a trend of increased expression (log2FC>1.8, p= 0.05 - 0.1).

In addition, miRNA expression was subsequently compared between day 0 and day 2. This analysis revealed significant downregulation of seven miRNAs with the largest decrease observed for hsa-miR-6743-3p, followed by hsa-miR-491-3p, hsa-miR-4746-5p, hsa-miR-1909-3p, hsa-miR-4745-3p, hsa-miR-6790-5p, and hsa-miR-6718-5p (Figure 2c). Notably, hsa-miR-6743-3p and hsa-miR-1909-3p that were increased during LOS subsequently decreased significantly by day 2. Moreover, hsa-miR-7108-5p, hsa-miR-3940-5p, and hsa-miR-6851-3p, which showed a suggestive increase during LOS, also tended toward decreased expression by day 2. Furthermore, no miRNAs were significantly differentially expressed between day 2 and control samples (Figure 2d).

To investigate the functional relevance of these findings, predicted target genes were examined by RT-qPCR in an independent cohort. *TMEM120B* was selected as a predicted target of hsa-miR-6743-3p, and *PTER* and *DRAM2* as predicted targets of hsa-miR-7108-5p. Paired day 0 and day 2 samples were available from five neonates (P11–P15; 10 samples) and were compared with two controls (C07–C08). *DRAM2* showed the most consistent pattern, with reduced expression in all five day 0 samples (mean log2FC = −1.67) and increased expression in four of the five samples obtained on day 2, approaching control levels (mean log2FC = −0.21, paired p=0.270). This pattern was inverse to the observed increase of hsa-miR-7108-5p during the LOS acute phase. *PTER* was also reduced in four of the five samples obtained on day 0 (mean log2FC = −0.96), and increased overall by day 2, although changes were not statistically significant (mean log2FC = 1.30; p=0.337). Similarly, *TMEM120B* was reduced in four of the five samples obtained on day 0 (mean log2FC = −0.56), with variable day 2 expression (mean log2FC = 0.70; p=0.433). Relative expression of *TMEM120B*, *PTER*, and *DRAM2* genes are shown in the Online Resource 5.

Finally, associations between normalized monocyte miRNA expression and systemic inflammatory indices (NLR and SII) were assessed within the NGS cohort (days 0 and 2), for each of the five miRNAs. At day 0 hsa-miR-7108-5p showed a significant positive correlation with NLR (ρ=0.659, p=0.038; Figure 3). No statistically significant correlation with NLR was observed for the remaining miRNAs (Figure 4). Additionally, none of the miRNAs correlated significantly with SII (data not shown).

**Fig. 3.**
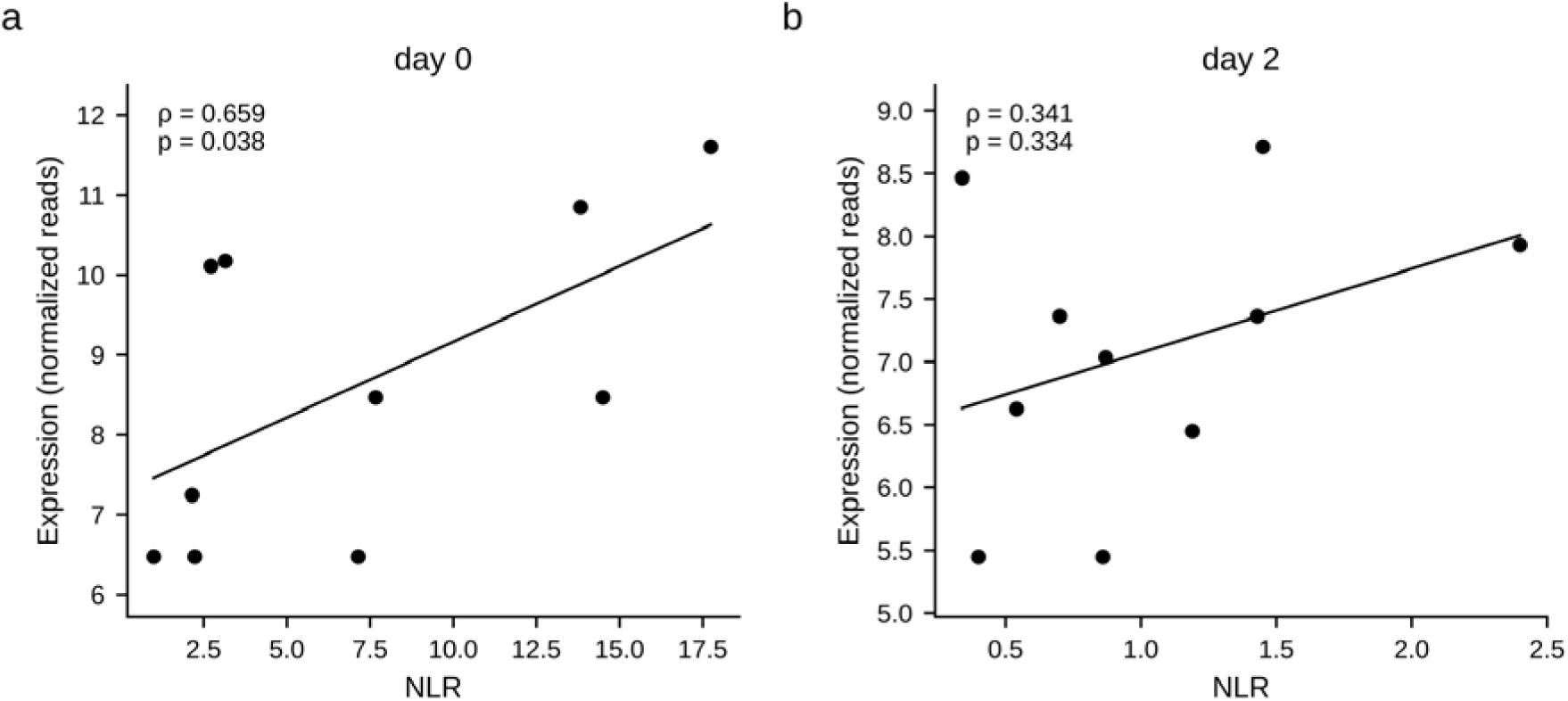
Correlation between hsa-miR-7108-5p expression and neutrophil-to-lymphocyte ratio (NLR) in neonates with late-onset sepsis (NGS cohort, n = 10). Each dot represents one neonate; the line shows a linear fit for illustration. Spearman’s rank correlation was assessed at (a) diagnosis (day 0; ρ = 0.659, p = 0.038) and (b) 48 hours post antibiotics (day 2; ρ = 0.341, p = 0.334)

**Fig. 4.**
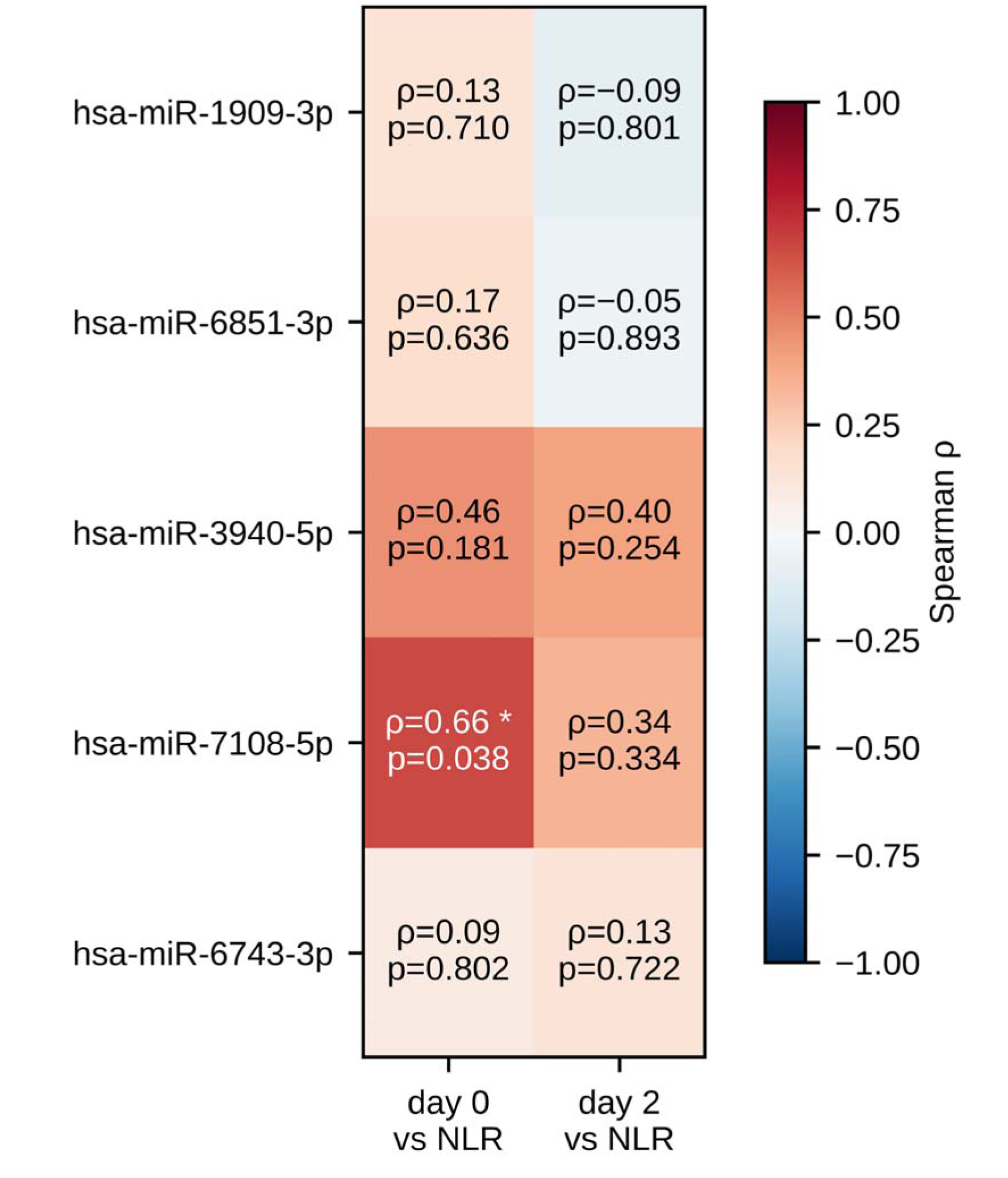
Correlation between miRNA expression levels and neutrophil-to-lymphocyte ratio (NLR) in neonates with late-onset sepsis (NGS cohort, n = 10). Heatmap showing Spearman’s rank correlation coefficients (ρ) and p-values at diagnosis (day 0) and 48 hours post antibiotics (day 2); the shading scale ranges from ρ = −1 to 1. The asterisk (*) indicates statistical significance (p < 0.05)

## Discussion

This study characterized monocyte-specific miRNA expression in neonatal LOS cases compared to controls, using NGS. Clinical severity was assessed using the complete blood count-derived NLR and SII. Across the total cohort, NLR was significantly elevated at LOS diagnosis, while both NLR and SII decreased by day 2. Hsa-miR-6743-3p was markedly upregulated during acute LOS, alongside increased expression of several other candidate miRNAs. These changes were transient, with miRNA expression approaching control levels by day 2, coinciding with clinical improvement following antibiotic treatment.

Few studies have used NGS for comprehensive miRNA discovery, particularly in monocytes or neonatal populations. In adult sepsis, small RNA sequencing of peripheral blood, including CD14^+^ monocytes, identified a distinct miRNA signature, different from that observed in our study, that distinguished sepsis from systemic inflammatory response syndrome with 86% accuracy and was validated by RT-qPCR in an independent cohort [18]. Similarly, combined miRNA and mRNA sequencing of peripheral blood monocytes identified distinct miRNAs and miRNA–mRNA regulatory networks, conceptually similar to our target-gene approach [19]. More recently, serum small RNA sequencing identified prognostic miRNA and tRNA-derived signatures associated with sepsis mortality [20], while serum exosomal sequencing followed by independent RT-qPCR validation identified miR-193b-5p as a diagnostic biomarker associated with CRP, PCT, and IL-6 [21]. Beyond infectious diseases, NGS-based discovery followed by RT-qPCR validation has also been applied in pediatric populations, including the identification of plasma extracellular vesicle-derived miRNA signatures in epilepsy [22]. Consistent with our findings, previous studies have not identified a common miRNA signature, suggesting that unbiased profiling may reveal context- or compartment-specific signals rather than universally dysregulated miRNAs. However, given the limited number of NGS-based studies, it remains unclear whether this lack of convergence reflects true biological heterogeneity or simply the limited available evidence.

Previous studies of neonatal LOS have mainly used targeted RT-qPCR of pre-selected serum/plasma miRNAs rather than unbiased profiling. Reported candidates include miR-34a-5p, miR-199a-3p [11]miR-16, miR-181a, and miR-23b, with altered expression, showing even better diagnostic performance than CRP and PCT (miR-16) [12].

A longitudinal study associating miR-15a and miR-223 with nSOFA scores [23] further supports a potential relationship between miRNA expression and disease severity. None of the miRNAs identified in our monocyte-specific NGS analysis—hsa-miR-6743-3p or the trend-level candidates hsa-miR-7108-5p, hsa-miR-3940-5p, hsa-miR-6851-3p and hsa-miR-1909-3p—overlapped with previously reported serum/plasma miRNAs. To our knowledge, these miRNAs have not previously been reported in the context of sepsis, infection, or immune regulation. This lack of overlap may reflect differences between monocyte and circulating miRNA profiles, although differences in study populations and methodology cannot be excluded. Previous studies in other diseases have suggested that cellular blood miRNA profiles may provide different or potentially more informative candidate biomarkers than circulating miRNAs [24].

Our findings add to evidence supporting NLR as a practical, low-cost marker for neonatal LOS [3,4]. Its discriminative performance in our cohort (AUC 0.958) was higher than that reported in a similar population (AUC 0.903) [4], although this comparison should be interpreted cautiously given our small sample size. In contrast, SII was not significantly associated with LOS in our cohort, differing from Zhu et al. [3], who identified SII as the strongest index evaluated among the indices studied. The observed positive correlation between hsa-miR-7108-5p and NLR at day 0 provides preliminary evidence of a link between molecular and clinical inflammatory markers, broadly consistent with findings in adult sepsis linking serum exosomal miR-193b-5p with CRP, PCT, and IL-6 [21]. From a clinical perspective, NLR may offer particular translational value, as it is derived from the routine complete blood count already included in the diagnostic evaluation of neonatal sepsis and requires no additional testing. Analysis of predicted miRNA targets provided further functional context. The inverse expression pattern of hsa-miR-7108-5p and *DRAM2* are compatible with a regulatory relationship during LOS and following treatment, whereas the variability of other targets highlights the complexity of miRNA-mediated regulation. These findings remain exploratory and require direct functional validation of the predicted miRNA–target interactions.

This study has several strengths. First, unbiased NGS profiling enabled the identification of miRNA signals beyond pre-selected candidates. Second, miRNAs were isolated specifically from monocytes, key innate immune cells involved in bacterial recognition and septic inflammation, rather than from unfractionated serum or plasma. Third, the longitudinal day 0/day 2 design captured treatment-related changes and enabled paired within-patient comparisons, raising the possibility that monocyte miRNA expression may reflect not only LOS diagnosis but also its clinical course. Fourth, NGS findings were evaluated in an independent RT-qPCR cohort and in relation to two low-cost clinical severity indices, providing complementary context for the discovery findings. Finally, despite the small sample size, prospective sampling before antibiotic administration represents an important strength, given the challenges of obtaining neonatal samples within this critical early window. Regarding study limitations, it should be noted at first, the small sample size may have limited the statistical power of the miRNA expression and clinical severity analyses. Secondly, miRNA profiling was performed in isolated monocytes rather than serum or plasma, limiting direct comparison with studies of circulating miRNAs. Moreover, the high discriminatory performance and cutoff identified for NLR were derived from the same small study population and may therefore overestimate performance in independent populations. Finally, monocyte miRNA expression may be influenced by gestational age, comorbidities, inflammatory status, medication exposure, and other clinical factors that could not be fully accounted for in this cohort. Larger multicenter studies with independent molecular and clinical validation are required before the identified miRNAs or NLR thresholds can be considered for clinical applications.

**In conclusion**, this study provides an unbiased characterization of the monocyte miRNA expression in neonates with LOS. Hsa-miR-6743-3p, was significantly altered at LOS diagnosis, while several additional miRNAs showed suggestive changes, with the overall expression pattern shifting toward baseline within 48 hours post antibiotics, alongside improvement in NLR. NLR, but not SII, showed high diagnostic performance within this cohort, supporting its potential as a simple, low-cost clinical marker. A positive association between hsa-miR-7108-5p and NLR was also observed, while its predicted target, *DRAM2*, showed an inverse expression pattern, with reduced expression during acute LOS and partial recovery following treatment. Although exploratory, these findings provide preliminary functional context for the identified miRNA signature and require further validation in larger multicenter cohorts before their clinical utility can be established.

## Acknowledgements

The authors gratefully acknowledge the participating families and the NICU staff for their invaluable contribution to this study.

## Author Contributions

EG: Investigation, Formal analysis, Writing – original draft, review and editing; AK: Investigation, Formal analysis, Writing – original draft, review and editing; EP: Investigation, Formal analysis, Writing – original draft, review and editing, Resources; MA: Formal analysis, Writing – review and editing; GD: Funding acquisition, Resources, Writing – review and editing; VS: Conceptualization, Project administration, Methodology, Formal analysis, Writing – review and editing, Supervision; DG: Conceptualization, Project administration, Methodology, Formal analysis, Writing – review and editing, Funding acquisition, Supervision

## List of Abbreviations

ACTB: Beta-actin (reference gene)
AUC: Area under the curve
BW: Birth weight
CPAP: Continuous positive airway pressure
CRP: C-reactive protein
DRAM2: DNA damage regulated autophagy modulator 2 (predicted target gene)
GA: Gestational age
GEO: Gene Expression Omnibus
HFNC: High-flow nasal cannula
IL-6: Interleukin-6
IQR: Interquartile range
LOS: Late-onset sepsis
miRNA(s): MicroRNA(s)
NGS: Next-generation sequencing
NICU: Neonatal intensive care unit
NLR: Neutrophil-to-lymphocyte ratio
nSOFA: Neonatal Sequential Organ Failure Assessment
PCT: Procalcitonin
PTER: Phosphotriesterase-related protein (predicted target gene)
ROC: Receiver operating characteristic
RT-qPCR: Quantitative real-time polymerase chain reaction
SII: Systemic immune-inflammation index
SNAP-II: Score for Neonatal Acute Physiology II
TMEM120B: Transmembrane protein 120B (predicted target gene)

## Statements and Declarations

### Data Availability

All data are available in anonymized form, with participants identified only by a unique study ID. No personally identifiable information is included in the dataset. The anonymized data are available from the corresponding author upon reasonable request.

### Ethics Approval

The study was conducted in accordance with the principles of the Declaration of Helsinki. Ethical approval was obtained from the Ethics Committee of the University General Hospital of Patras (Number 28049/03/11/21).

### Consent to Participate

Informed consent was signed by both parents prior to enrollment for both cases and controls.

### Consent for Publication

Not applicable.

## Supplementary Material

**Online Resource 1.** Clinical LOS criteria*.

| Clinical LOS criteria |
| --- |
| Temperature instability |
| Apnea or increased oxygen requirements |
| Poor peripheral perfusion or hypotension |
| Bradycardia or tachycardia |
| Feeding intolerance, abdominal distension |
| Lethargy, irritability |

\*European Medicines Agency. Report on the expert meeting on neonatal and paediatric sepsis. London: European Medicines Agency; 2010.
https://www.ema.europa.eu/en/documents/report/report-expert-meeting-neonatal-paediatric-sepsis_en.pdf

**Online Resource 2.**
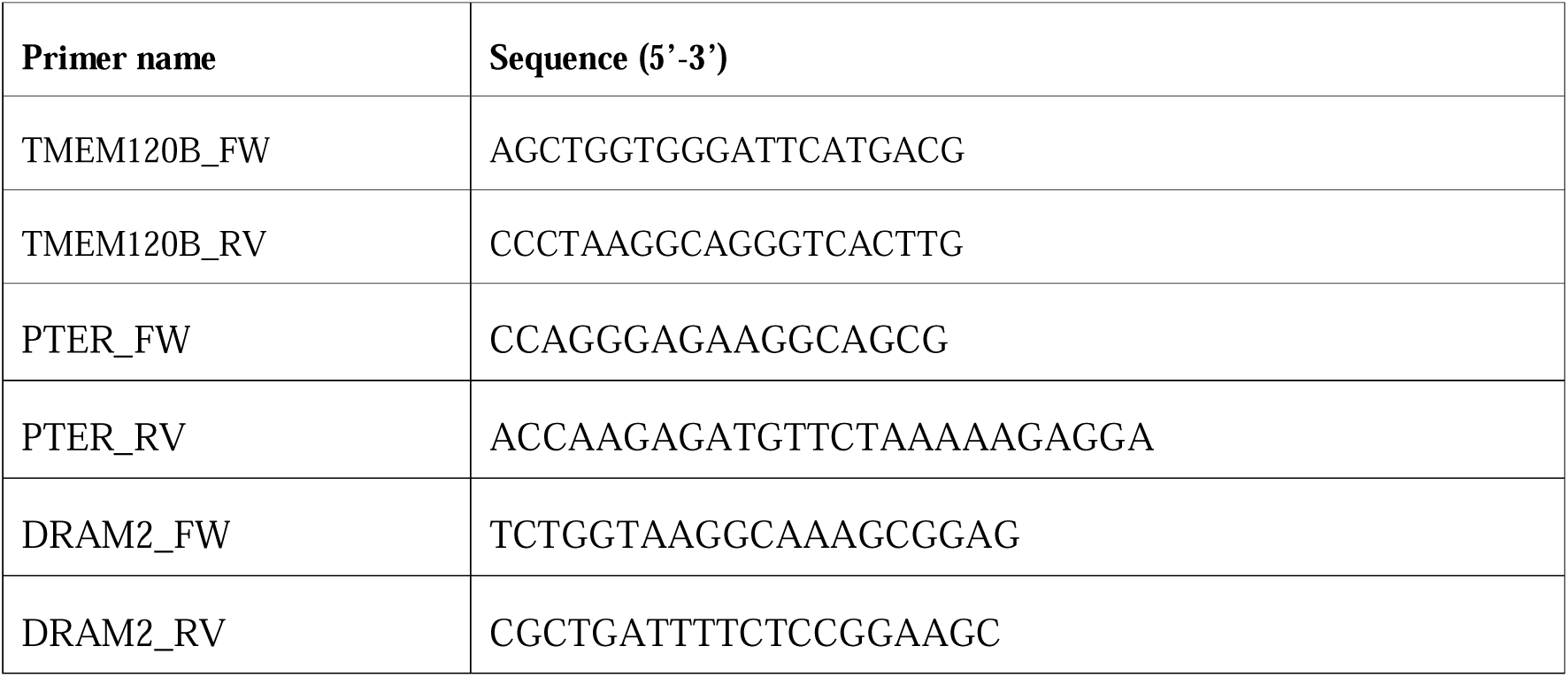
RT-qPCR primer sequences for target gene expression analysis.

**Online Resource 3.**
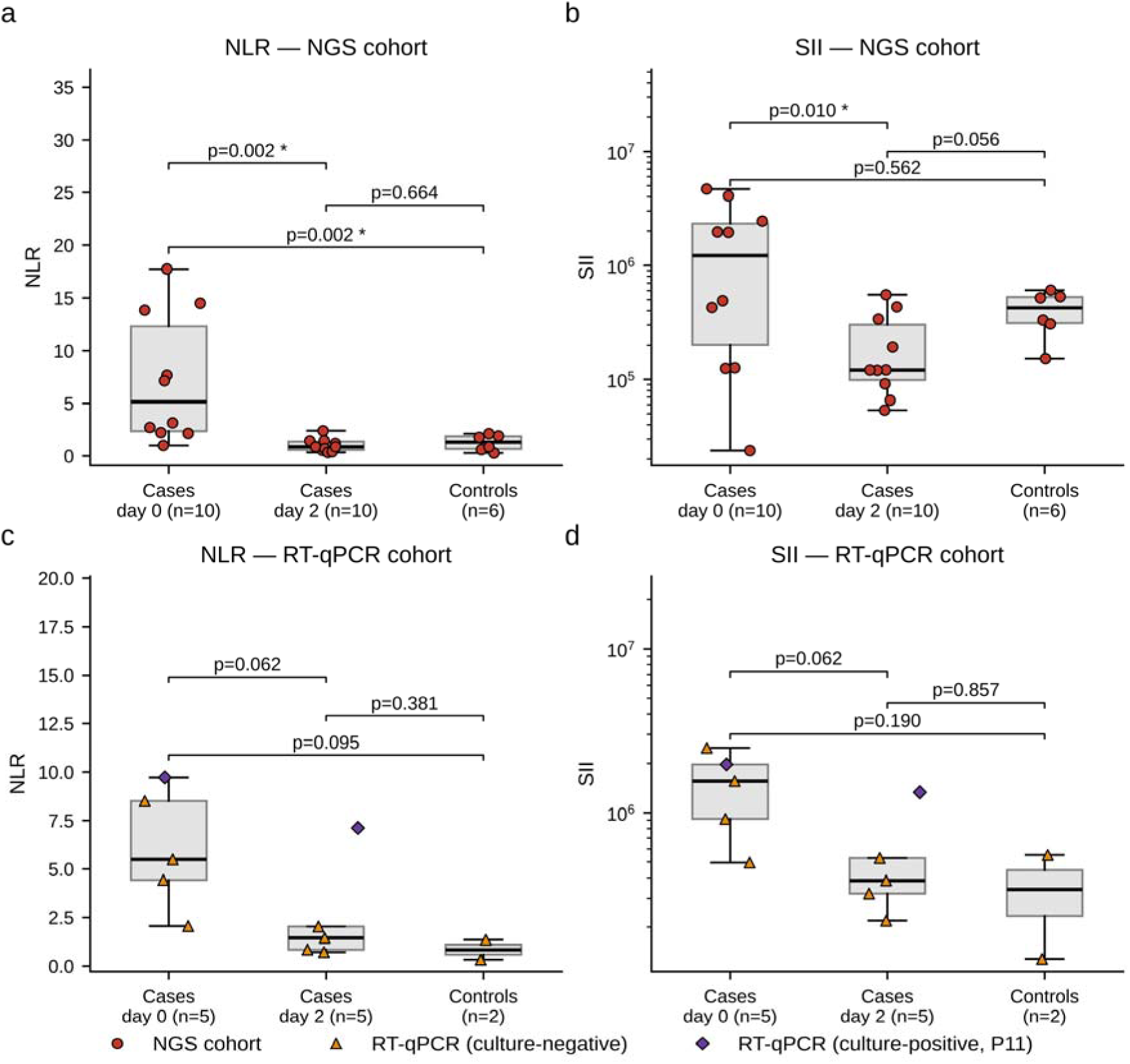
Systemic inflammatory indices (NLR, SII) by cohort: NGS cohort (a, b, LOS cases, n = 10 vs healthy controls, n = 6) and RT-qPCR cohort (c, d, LOS cases, n=5 vs healthy controls, n=2)

**Online Resource 4.**
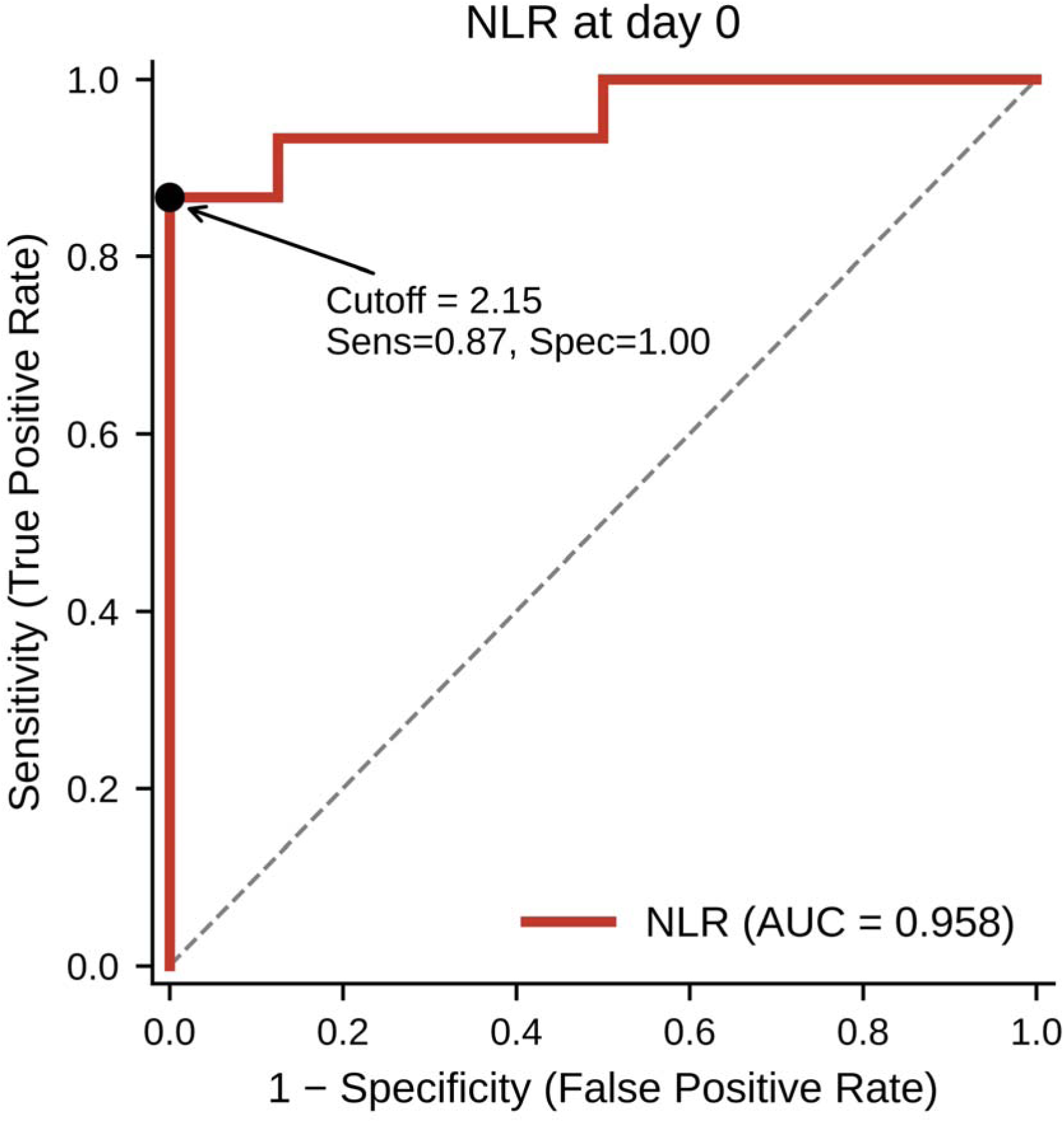
ROC curve for discrimination of LOS cases vs healthy controls (n=15 cases, n=8 controls)

**Online Resource 5.**
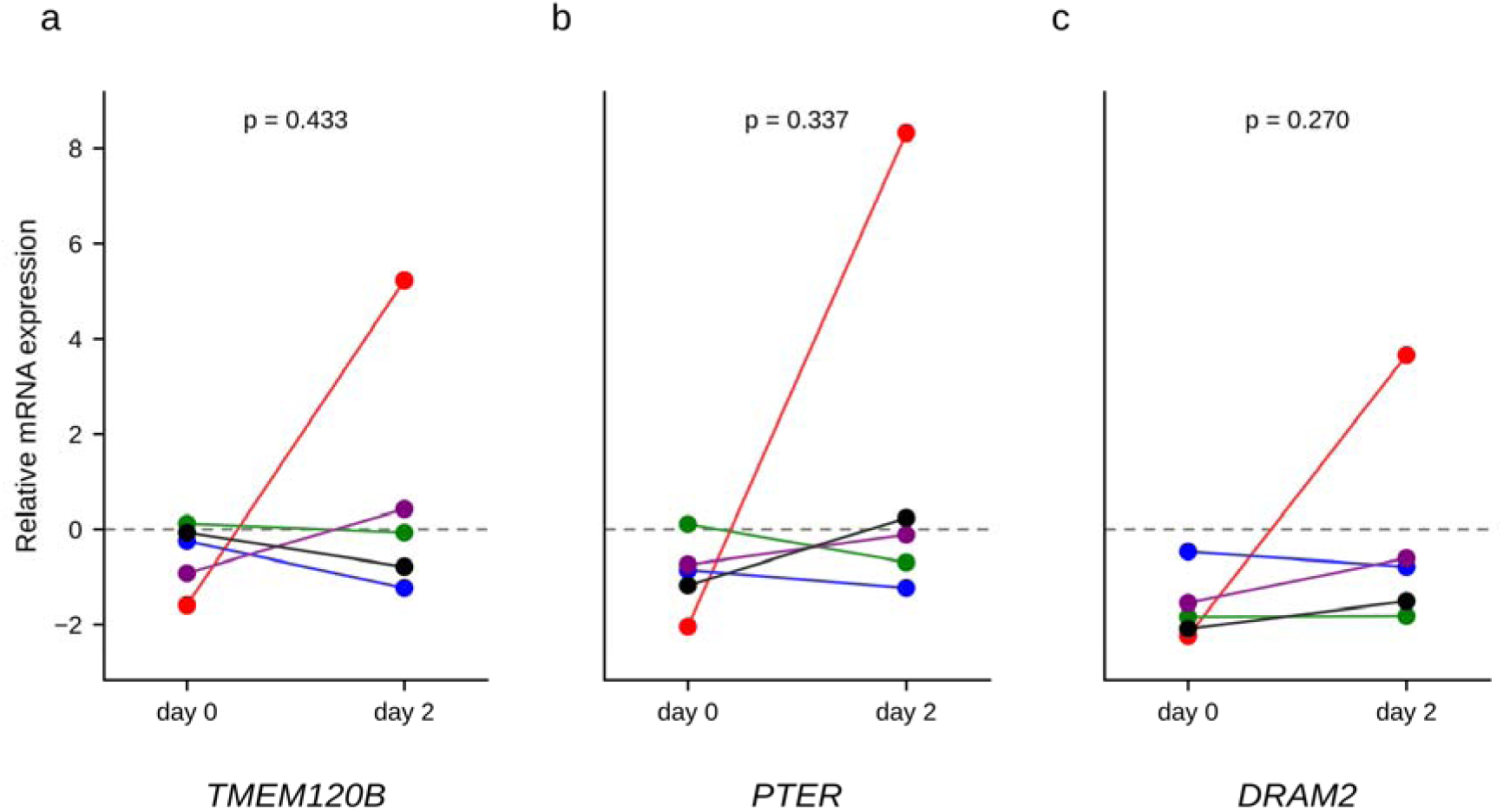
Expression analysis of predicted miRNA target genes at day 0 and day 2. Relative mRNA expression levels of *TMEM120B* (a), *PTER* (b), and *DRAM2* (c) were determined by RT-qPCR in paired samples collected from neonates with late-onset sepsis (LOS) at day 0 and day 2 (n=5). Expression levels are presented relative to the healthy control group, with the dashed horizontal line at 0 representing the control reference level. Each color represents an individual patient, and paired measurements from the same patient on day 0 and day 2 are connected by lines. Statistical comparisons between day 0 and day 2 were performed using a paired t-test

